# Implicit processes do not contribute to learning to reach in small mirror reversed environments

**DOI:** 10.1101/2025.09.17.676913

**Authors:** Sarvenaz Heirani Moghaddam, Erin Krista Cressman, Gerome Aleandro Manson

## Abstract

Learning to reach with a small visuomotor rotation (VR; a rotation of visual feedback relative to hand motion) has been shown to arise unconsciously (i.e., implicitly). Whether the same processes support learning in a small mirror reversal (MR), where feedback is reflected across the body midline, remains unknown. To address this gap, we asked whether implicit processes contribute to learning in a small MR. Forty-two right-handed participants reached to targets located 10° to the left and right of body midline using a Kinarm exoskeleton robot. Half of the participants experienced a VR distortion (VR group), which consisted of a 20° clockwise or counterclockwise cursor rotation. The remaining participants experienced a 20° MR distortion (MR group), where cursor feedback was reflected across body midline (y-axis). Following reaches with a VR or MR distortion, participants completed assessment trials in which they reached in the absence of cursor feedback to assess implicit learning. Analysis of angular errors (AE) revealed that all participants in the VR group learned to reach with the VR distortion, however, only 55% of MR participants learned to reach with the MR distortion. AEs on the no-cursor trials revealed that only the VR group engaged in implicit learning. These findings demonstrate that MR learning, even when small MR distortions are introduced, is not supported by implicit learning. The absence of implicit learning in MR provides evidence that MR is a different form of learning (i.e., skill acquisition) compared to VR learning (i.e., motor adaptation).

## Introduction

Motor learning is a broad term encompassing a range of concepts and experimental approaches (1,2). Traditionally, motor learning has been defined as a set of processes that lead to a relatively permanent change in behaviour as a result of practice (2). In 2019, Krakauer and colleagues broadened the scope of this definition to include both skill acquisition and skill maintenance under the umbrella term of motor learning. Skill acquisition refers to “the processes by which an individual acquires the ability to rapidly identify, select, and execute an accurate movement given a sensory stimulus and/or the current state of the body and world,” while skill maintenance, also commonly referred to as motor adaptation, refers to “the ability to maintain performance levels of existing skills under changing conditions.”

Motor learning has been studied in the laboratory by having participants reach in a novel visuomotor environment. For example, a cursor’s trajectory on the screen can be rotated 20° clockwise (CW, or rightwards) relative to hand motion (visuomotor rotation (*VR)*; (3–5). If a participant reaches directly to the target, the cursor veers slightly to the right of the target. In VR paradigms the magnitude of the cursor rotation is typically kept constant across all target locations, ensuring that the rotation experienced would be a 20° CW rotation relative to hand motion regardless of target location. Thus, a participant must reach 20° counterclockwise (CCW) or left of the target to get the cursor on the target.

Another example of altered visual feedback is mirror reversed (MR) cursor feedback (see Telgen et al., 2014b; Wang & Taylor, 2021b; Wilterson & Taylor, 2021). In a MR paradigm, the cursor motion on the screen is mirrored across body midline (i.e., y-axis) relative to hand motion. Thus, contrary to VR, the size and direction of the distortion between the cursor on the screen and hand motion is dependent on a participant’s hand position relative to body midline. If the target is placed 10° to the right of body midline, reaching directly to the target brings the cursor 10° to the left of body midline. In this specific example, the magnitude of the MR distortion remains constant at 20° when reaching directly to the right target. However, because cursor motion is mirrored across body midline, any deviation in hand position will result in a change in the distortion magnitude. For example, if a participant reaches 15° to the left of body midline, cursor feedback would be displayed 15° to the right of body midline, introducing a 30° distortion between cursor and hand motion. In MR paradigms, the direction of the distortion also changes from CW to CCW, depending on whether the participant is reaching to the left or right of body midline respectively. Reaching in a VR versus an MR environment has been shown to lead to differences in ‘aftereffects’. If participants are instructed to reach directly to the target in the absence of cursor feedback after reaching in a VR environment, they continue to reach as if the VR is still present (9–11). These aftereffects in VR learning are proposed to arise implicitly and reflect non-intentional changes to an existing internal model (i.e., sensorimotor mapping between hand motion and visual feedback) that arise due to experiencing a sensory-prediction error (12). Implicit processes have been shown to play the dominant role when one learns to reach with a small VR distortion (i.e., a cursor rotation less than 30°; see Modchalingam et al., 2019; Neville & Cressman, 2018; Werner et al., 2015).

In contrast to VR learning, implicit processes have not been shown to have a role in learning a large MR. For example, Wang and Taylor (2021) found no evidence of aftereffects immediately following reaches in an MR environment. Wilterson and Taylor (2021) also observed no aftereffects following 5 days of reaching in an MR environment, and Telgen et al. (2014) showed that reaching with an MR distortion produced no aftereffects on post-test trials that had aligned cursor feedback. One important caveat to these previous MR findings is that, in these studies, participants reached with large MR distortions (e.g., 90° MR: Wang and Taylor, 2021; 45° MR: Wilterson and Taylor, 2021; 40° MR: Telgen et al., 2014). These large distortions have been shown to engage explicit (i.e., conscious strategy) processes even in VR learning paradigms (13–15). For example, Neville & Cressman directly compared the contribution of implicit and explicit processes in learning of 20°, 40° and 60° VR and found that explicit processes only contributed to learning of a 40° and 60° rotations and not the 20° VR (2018). The role of implicit processes in learning a small MR has yet to be established.

As such, the goal of the current research was to determine the contributions of implicit processes to learning a small (20°) MR distortion. Specifically, we aimed to determine whether implicit processes, which are prominent in VR learning, also play a dominant role in the learning of a small MR distortion. Furthermore, we compared the contributions of implicit processes to learning a small MR versus a small VR distortion to gain insight into the forms of motor learning engaged. Telgen and colleagues (2014) have recently suggested that reaching with a VR distortion implicitly updates an existing sensorimotor mapping between hand motion and visual feedback and hence is a form of motor adaptation (i.e., skill maintenance), while reaching with a MR distortion leads to the acquisition of a new sensorimotor mapping (i.e., skill acquisition or de novo motor learning; see also Gastrock et al., 2024). We hypothesized that, contrary to learning a small VR distortion, implicit processes would have a negligible contribution to learning a small MR distortion. Thus, we expect that the MR group would show minimal aftereffects in the no-cursor trials after learning. The absence of implicit contributions to MR learning would suggest that MR learning is a distinct form of learning (e.g., skill acquisition) compared to VR learning (e.g., motor adaptation).

## Methods

### Participants

Forty-two participants (F = 25), aged 19-45 years (24.0 years ± 4.9 years) were recruited from the Queen’s University community and randomly divided into two groups; (1) VR group (N = 22; F = 13) and (2) MR group (N = 20; F = 10). The VR group was further divided into two subgroups of participants, including a VR counterclockwise group (CCW; VR-CCW group; N = 11; F = 6) and a VR clockwise group (CW; VR-CW group; N = 11; F = 8). Participants were naïve to the purpose of the experiment and had never participated in research involving altered visual feedback. The experimental protocol was approved by the Queen’s University Health Sciences and Affiliated Teaching Hospitals Research Ethics Board (HSREB) and the University of Ottawa Health Sciences and Science Research Ethics Board. The recruitment period for this study started on 22/01/2024 and ended on 09/07/2024.

Upon arrival at the laboratory, participants were provided written informed consent. Participants then completed the Edinburgh Handedness Inventory (M = 93, SD = 11), and all were determined to be right-handed. Participants also completed a brief neurological questionnaire (adapted from Miles, 1930), to ensure that they did not demonstrate neurological impairment. Participants were then provided with an overview of the experiment using a PowerPoint presentation.

### Experimental apparatus

Participants reached to targets using the Kinarm Exoskeleton Lab (Kinarm, Kingston, ON, Canada; Kasuga et al., 2022; Scott, 1999; Singh & Scott, 2003). The Kinarm was positioned adjacent to the experimenter’s computer workstation and consisted of a downward-facing computer monitor with a refresh rate of 120 hertz (Hz) and a reflective surface below the computer monitor (see Figure 1A for experimental setup). The downward facing monitor projected visual stimuli onto the reflective surface.

**Figure 1.**
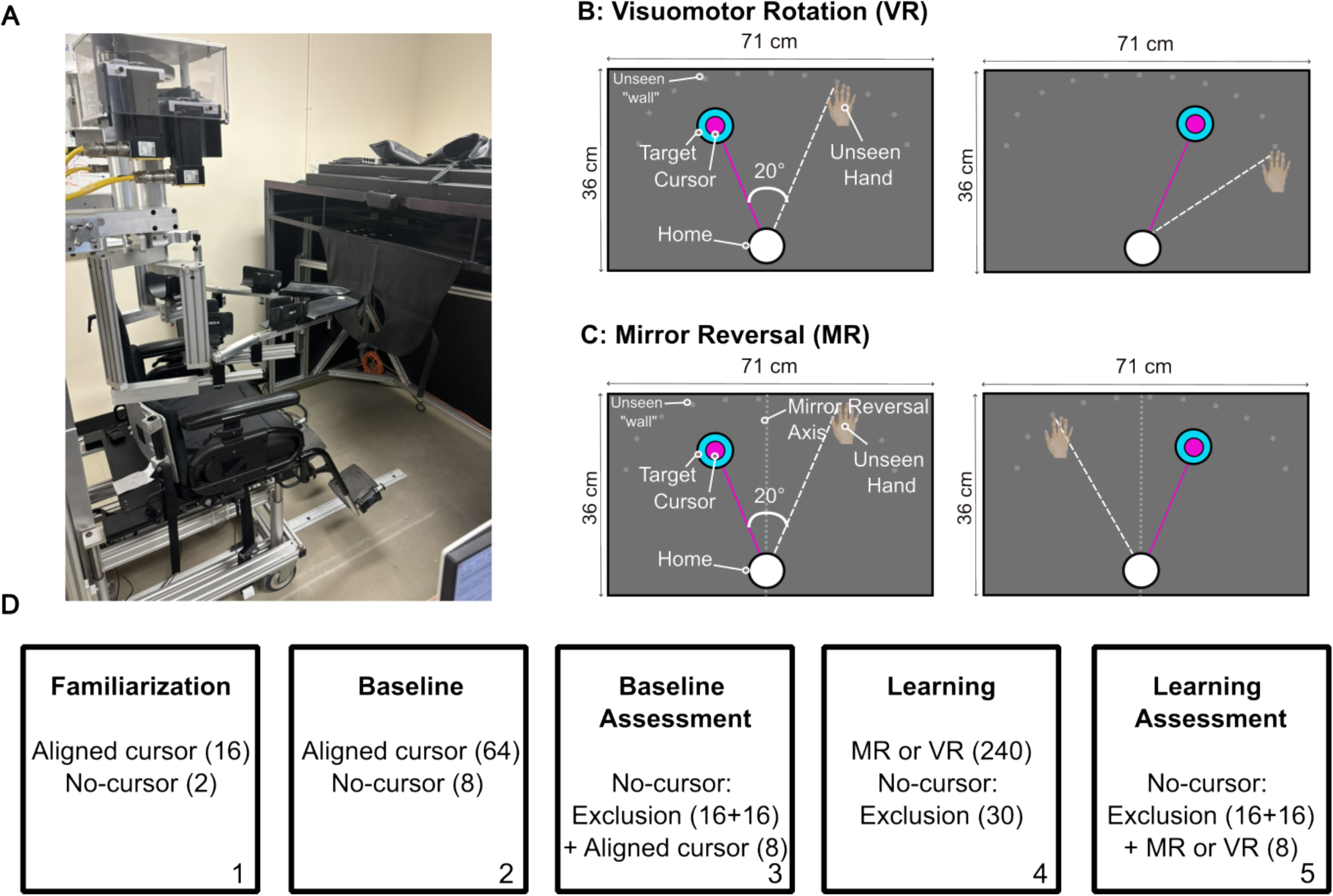
**A.** Experimental apparatus. Participants were seated in a wheelchair and reached to targets using the Kinarm Exoskeleton robot. **B and C.** Visual stimuli displayed during reaching trials with the visuomotor distortions. Target placement for this experiment included two targets positioned 10° to the right and left of center (i.e., 20° apart) at a distance of 10 cm from the home position. **B**. For the visomotor rotation (VR) groups, the cursor motion was rotated 20° clockwise (CW) or counterclockwise (CCW) relative to their index finger motion. **C.** For the mirror reversed (MR) group, the cursor motion was mirrored across the y-axis relative to index finger motion. When the left target was shown, the index finger needed to move 20° to the right of the target in order to get the cursor on the target and when the right target was shown, the index finger needed to move 20° to the left of the target in order to get the cursor on the target. The dotted line in the middle depicts the mirror axis, which was not visible to participants. **D**. Overview of blocks of trials and the number of trials (in parantheses) within each block completed by participants in each group.

Participants were seated in a height-adjustable wheelchair, and the experimenter adjusted the height and distance of the chair from the reflective surface, such that the participant was able to comfortably see and reach the visual stimuli on the screen. Participants’ right and left arms were each placed in two size-adjustable troughs (one between the wrist and the elbow, and the other between the elbow and the shoulder). The left arm remained stationary throughout the experiment. Participants moved their right arm through flexion, extension, abduction, and adduction of the elbow and the shoulder in the horizontal plane. The position of the right index finger was represented on the screen as a cursor (i.e., white circle, 0.5 cm in diameter) and the position of this finger was recorded at a sampling rate of 1000 Hz. The reflective surface and a velcro-secured drape around the participant’s neck blocked their vision of their right limb.

### Types of Trials

Figure 1D provides an overview of the block structure within the experiment. The blocks of trials included familiarization, baseline, baseline assessment, learning and learning assessment. A detailed description of the trial types completed in each block is provided in the next section and illustrated in Figure 2.

**Figure 2.**
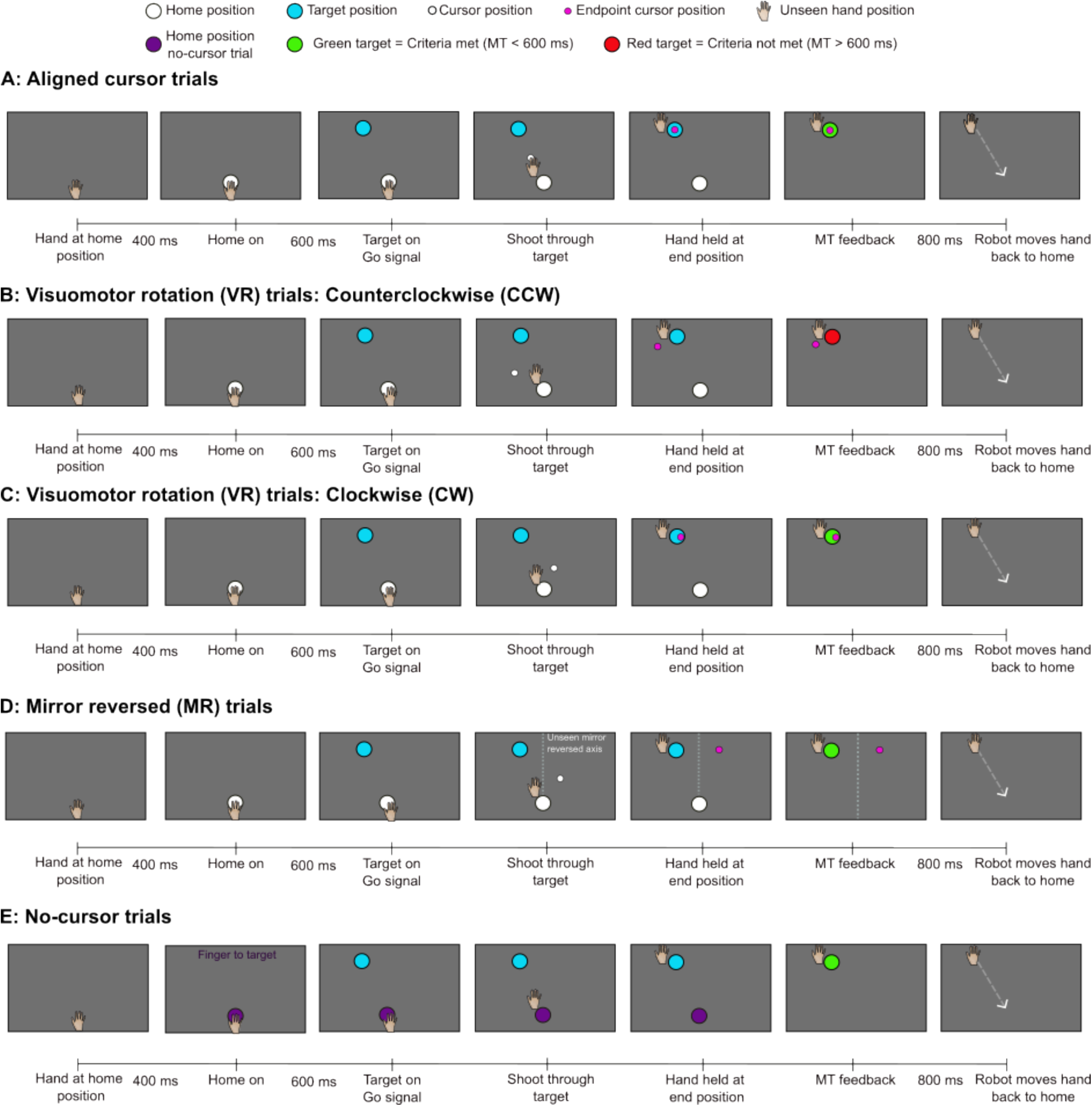
Trials completed and accompanying timeline. The hand and index finger were hidden from participants’ view. **A.** Aligned cursor trials: Reaches with aligned cursor feedback. The cursor was aligned to a participant’s index finger position and was continuously visible throughout the reach until the finger crossed the target array. **B-C.** VR trials: The cursor’s trajectory was rotated 20° CCW (**B**) or 20° CW (**C**) relative to the trajectory of the index finger and was visible throughout the reach until the finger crossed the target array. **D.** MR trials: Reaches with mirror reversed feedback. The cursor’s trajectory was mirrored across the y-axis relative to the trajectory of the index finger and was visible throughout the reach until the finger crossed the target array. **E.** No-Cursor trials: Participants were instructed to reach to the target. Movement time (MT) feedback was provided at the end of each shooting movement, before the robot moved the hand back to the home position to begin the next trial.

### Aligned cursor trials

Participants made reaching movements towards two targets that were 10 cm from the home position. The targets were blue circles (0.75 cm in diameter) that appeared 10° to the right and left of the y-axis, which was aligned with a participant’s midline (Figure 1B). The two targets appeared with equal probability. In aligned cursor trials, participants reached while a white cursor (0.5 cm in diameter) accurately represented their index finger position on the screen (Figure 2A).

Each trial began with the participants holding their index finger at the home position for 400 ms. The home position and the cursor then appeared. Following another 600 ms, one target appeared. Participants were instructed to move the cursor through the target as quickly and accurately as possible by performing a slicing (or shooting) action. This slicing action was completed as a ballistic shooting movement such that a participant’s index finger passed through the target at approximately peak velocity (20). Movements were terminated by a soft wall placed 2 cm beyond the target. This soft wall (spring constant 150 N/m) made the movement feel more natural and encouraged participants to move rapidly through the target. Visual feedback regarding the right index finger position was available up until the participant’s index finger crossed the target array. At this time, the cursor stopped, and participants no longer received visual feedback about their index finger’s position. Movement onset was defined online as the time when the centre of the cursor moved 0.5 cm away from the centre of the home position and when velocity first increased above 0.03 m/s and remained above 0.03 m/s for the subsequent 9 ms. Movement end was defined as the time that the hand crossed the target array. Movement time (MT) was defined as the time between movement onsent and movement end.

Participants received visual feedback regarding their MT in the form of a target colour change once their index finger reached the soft wall. The goal MT was less than 600 ms, such that the target turned green if MT was less than 600 ms and red if MT was greater than 600 ms. On all trials, the hand remained in the movement end location for 800 ms before the robot moved the participant’s index finger back to the home position along a linear path in an MT of 1000 ms in the absence of cursor feedback to start the next trial (see Figure 2). If participants attempted to move out of the linear path, an equal and opposite force (3.3 newton meters, Nm) was applied to their hand to keep the hand within the linear path.

### VR and MR trials

These trials were similar to aligned cursor trials but participants experienced a VR (Figure 2B & 2C; VR group) or MR (Figure 2D; MR group) distortion. For the VR groups, visual feedback of the index finger position was rotated 20° CW (VR-CW group) or CCW (VR-CCW group) relative to hand motion. For the MR group, visual feedback of the hand position was mirrored across the y-axis. As a result, reaching directly to the right target (10° right of midline) caused the cursor to appear 10° left of midline, producing a 20° distortion between the cursor and hand motion.

### No-cursor trials: Exclusion trials

No-cursor trials were similar to aligned cursor trials except that the cursor remained off during the reach (Figure 2E). These trials were used to establish implicit processes (i.e, aftereffects) in accordance with the Process Dissociation Procedure (PDP; adapted from Werner et al., 2015). At the beginning of the exclusion trials, participants were instructed:

> “You are now going to reach when you cannot see your index finger, as there will be no-cursor on the screen. Do not use anything you may have learned to get the cursor to the target. Instead, aim so that your index finger goes straight through the target as you did during baseline reaches.”

The absence of the cursor in no-cursor trials was cued with a change in colour of the home position, such that on these trials the home position colour was purple and the instruction “Finger to target” appeared on the screen above the target.

In the familiarization, baseline and learning blocks, one no-cursor trial was completed for every 8 cursor trials. In the baseline block, participants completed 64 aligned cursor reaches and 8 no-cursor reaches. In the learning block, participants reached with either a VR or MR cursor distortion for 240 trials and completed 30 no-cursor trials. In the baseline and learning assessment blocks, participants completed 16 no-cursor exclusion trials, followed by 8 reaching trials with cursor feedback and then an additional 16 no-cursor exclusion trials. The 8 cursor trials were included to ensure that any learning was maintained.

### Data Analyses

All reaching trials were analyzed using custom-written MATLAB scripts (MATLAB R2022a, The MathWorks, Inc.). Hand angular error at the movement endpoint (AE), defined as the angular difference between a reference vector connecting the home position to the target and a reaching vector connecting the home position to the finger position at the end of the movement, was the main variable of interest. RT and MT were also analyzed. RT was defined as the time between target onset and movement start (defined as the time when the centre of the cursor moved 0.5 cm away from the centre of the home position and when velocity first increased above 0.03 m/s and remained above 0.03 m/s for the subsequent 9 ms). MT was defined as the time from movement start until movement end (defined as the time that the index finger crossed the target array).

### Outlier procedures

The following variables were used to screen for outliers: Start X and Start Y (x and y coordinates of the movement’s start position), AE and MT. If any of these variables was greater than 3 standard deviations (SD) from the participant’s average on a specific type of trial within each block (baseline, baseline assessment, learning, or learning assessment), the trial was removed from further analyses. For example, an aligned cursor trial was compared to the mean AE of the aligned cursor reaches within that block, while a no-cursor trial was compared to the mean of the no-cursor trials within the same block. Any reach with an MT greater than 600 ms or below 100 ms was flagged as an outlier and removed from analyses. Across all groups, participants completed 14,226 trials. 926 (6.51%) of these trials were classified as outliers and removed from the analyses presented below.

### Learning

Reaches with cursor feedback in the baseline and learning blocks were designated as early and late trials. The first 24 trials were designated as early trials and the last 24 trials as late trials. In order to assess implicit contributions to motor learning, we required participants to learn the VR or MR distortion. Thus, each participant’s data were screened to determine if they demonstrated learning by examining individual learning curves and comparing a participant’s average AE in the late learning phase (last 24 trials) with their AE in the late baseline phase (last 24 trials) for both right and left targets. To confirm learning, participants had to show a difference in AE in the late learning phase compared to the late baseline phase that was greater than 3 times the standard deviation of reaches in the late baseline phase. Only participants who met this learning criterion were included in subsequent analyses. Using this criterion, we found that all participants in the VR groups learned, while only 11 participants in the MR group demonstrated learning. Data related to the remaining 9 non-learning participants in the MR group are included in the Supplementary File. For all participants who learned, early and late AE data for each target in the learning block were nomalized by subtracting early and late AE in the baseline block.

The absolute value of these normalized AE were then compared between groups in a 3 group (MR, VR-CW, VR-CCW) x 2 target (right, left) x 2 time (early trials, late trials) mixed analysis of variance (ANOVA) with repeated measures (RM) on the last two factors. We also compared AE variability and RT across groups in a 3 group (MR, VR-CW, VR-CCW) x 2 target (right, left) x 2 time (early trials, late trials) mixed ANOVA with RM on the last two factors. To confirm that participants across all groups complied with the MT criteria, MT was analyzed in a 3 (MR, VR-CW, VR-CCW) × 2 target (right, left) × 2 time (early trials, late trials) mixed ANOVA with RM on the latter two factors.

### Implicit Learning

An implicit index was calculated for each target within the baseline and learning assessment blocks. The implicit index was defined as the mean AE of the no-cursor trials:

1. Implicit index = *x^-^*_AE of no-cursor trials_ These indices were then used to calculate implicit learning according to the following formula:
2. Implicit learning = Implicit index (learning assessment) – Implicit index (baseline assessment)

To allow for meaningful comparisons across groups and targets, all AE values were directionally normalized so that positive values reflected movement in the expected direction of learning, regardless of target.

Implicit learning for the right and left targets were compared across groups in a 3 group (MR, VR-CW, VR-CCW) x 2 target (right, left) mixed ANOVA with RM on the last factor. We also compared the magnitude of implicit learning to zero for each group by performing a Bayesian one-sample t-test. These t-tests allowed us to establish the presence of implicit learning following MR (and VR) learning.

All analyses were completed using JASP, R and Microsoft excel softwares. In the case where the test of homogeneity of variance (Levene’s test) was violated, a Welch correction was performed. Further, if the assumption of sphericity was violated, Greenhouse-Geisser corrected degrees of freedom were reported. The significance threshold for all statistical tests was set at *p* < 0.05 and post hoc tests with Bonferonni corrections for multiple comparisons were used to find the locus of significant effects or interactions. For clarity, only the breakdown of the highest-order significant interaction is reported.

## Results

### Learning

Mean AE in the baseline block was close to zero for all groups at all times (VR-CW group: early M = −0.3°, SD = 1.0°, late M = 0.1°, SD = 1.2°; VR-CCW group: early M = 0.2°, SD = 1.3°, late M = 0.4°, SD = 1.1°; MR group: early M = −0.2°, SD = 0.9°, late M = −0.2°, SD = 1.3°). AE did not differ significantly different between groups (*F*(2,30) = 0.760, *p* = 0.477, *η²* = 0.022) or target (*F*(1,30) = 1.061, *p* = 0.311, *η²* = 0.014).

Mean RT in the baseline block was not different between groups (*F*(2,30) = 0.727, *p* = 0.492, *η²* = 0.034). Analysis of mean RT in the learning block revealed a main effect of group (*F*(2,30) = 7.475, *p* = 0.002, *η²* = 0.208), indicating that the MR group (M = 287.8 ms, SD = 64.2 ms) showed significantly longer RT compared to the VR-CW group (M = 234.0 ms, SD = 49.4 ms; *p* = 0.006) and VR-CCW group (M = 232.0 ms, SD = 13.3 ms; *p* = 0.008). RT for the VR-CW group was not different from the VR-CCW group (*p* = 1.000).

Further, analysis of MT data confirmed that all participants adhered to the MT criteria, completing their reaches within 600 ms. ANOVA revealed no significant differences between groups (F(2,30) = 1.672, p = 0.205, η² = 0.060). However, there was a significant main effect of target (F(2,30) = 6.471, p = 0.016, η² = 0.011), with post hoc analyses revealing that reaches to the right target were approximately 11.6 ms faster than those to the left target. These results confirm that participants executed their movements consistently across groups and within the prescribed timing constraints.

With respect to learning, ANOVA revealed significant main effects of group (*F*(2,30) = 3.992, *p* = 0.029, *η²* = 0.043) and time (*F*(1,2) = 58.200, *p* < 0.001, *η²* = 0.247), as well as a significant group x time interaction (*F*(2,30) = 9.521, *p* < 0.001, *η²* = 0.081). Post hoc analysis of the group x time interaction indicated that early AE for the MR group was significantly less than AE for both the VR-CW group (*p* = 0.050) and the VR-CCW group (*p* = 0.022; Figure 3A and 3B). Additionally, AE in the early learning trials was significantly lower than the late learning trials for the MR group (*p* < 0.001; Figure 3B), whereas there were no reliable difference in AEs between the early and late learning trials for the VR-CW group (*p* = 0.185) or VR-CCW group (*p* = 0.223). The magnitude of AE in the late learning trials was not different across groups (MR group vs. VR-CCW group: *p* = 0.49, MR group vs. VR-CW group: *p* = 0.15, VR-CCW group vs. VR-CW group: *p* = 0.16). The main effect of target was not significant (*F*(1,30) = 3.569, *p* = 0.069, *η²* = 0.014).

**Figure 3.**
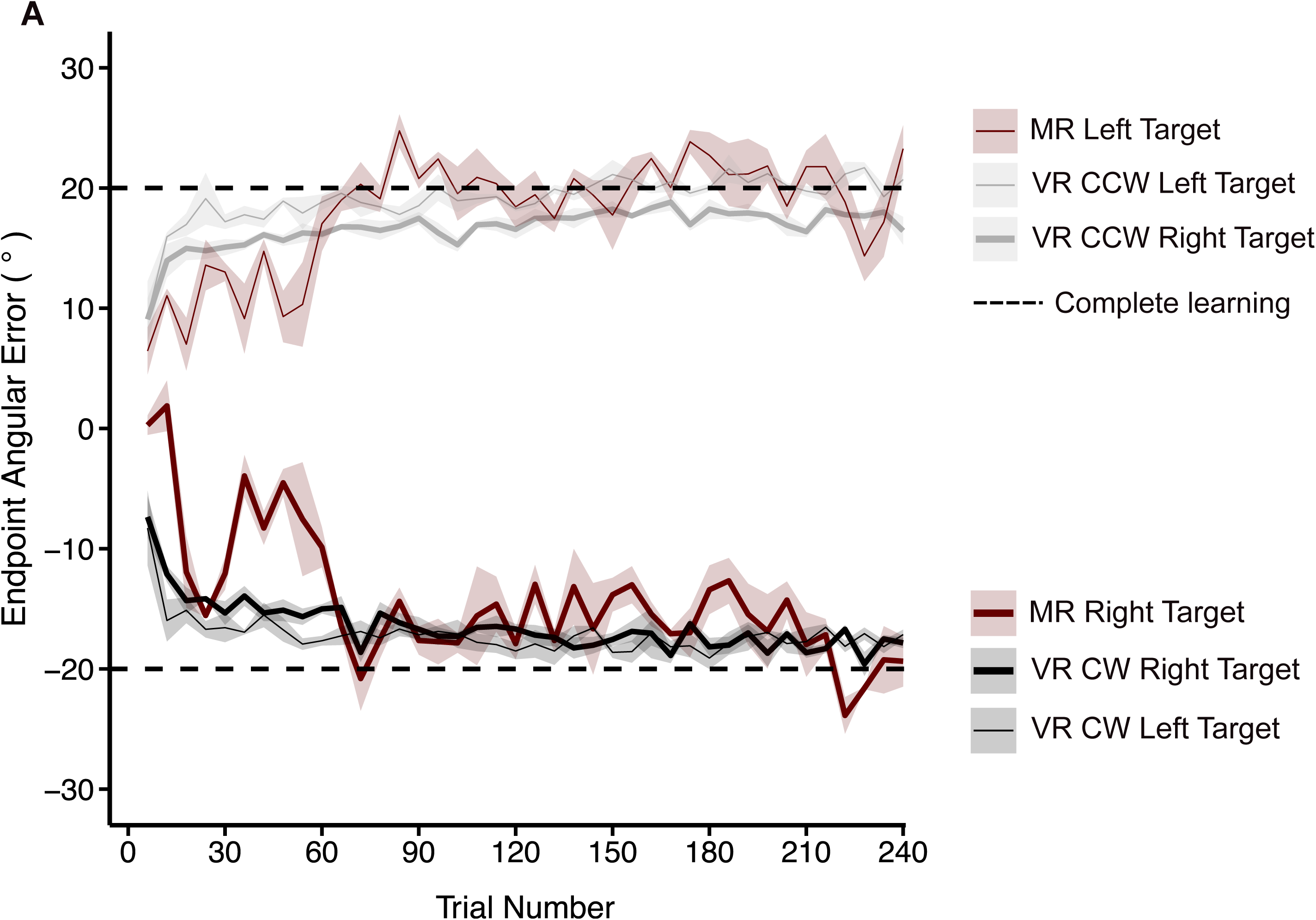
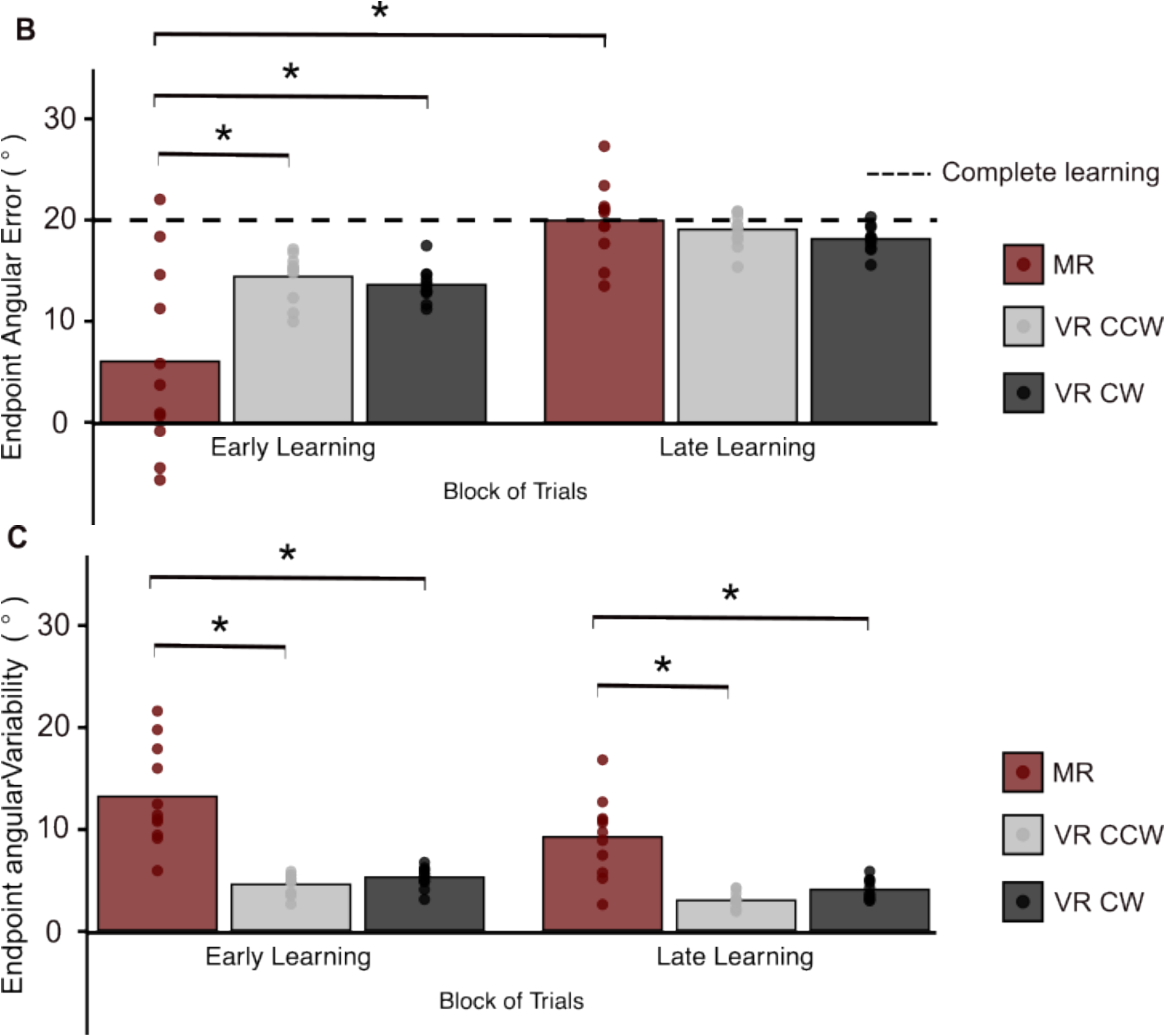
**A**: Learning curves for each group across trials. Endpoint angular errors (AE) in degrees are averaged over three consecutive trials in the learning block. Solid bold lines represent reaches toward the right target, while solid thin lines indicate reaches toward the left target. The shaded areas represent the standard error of the mean. Dashed lines mark the expected reach adjustments to demonstrate complete learning, with the dashed line at y = −20 showing complete learning for the right target and the dashed line at y = 20 showing complete learning for the left target. The MR group is depicted in maroon, the VR-CCW group is depicted in gray, and the VR-CW group is depicted in black. **B**: Absolute values corresponding to average AE during early and late learning for each group.The dashed line indicates complete learning. **C**: Average variability of AEs during early and late learning for each group. Maroon depicts the MR group, grey depicts the VR-CCW group and black depicts the VR-CW group. Individual participant data is shown by the circles. Asterisks denote group comparisons where differences were statistically significant (*p* < 0.05).

**Figure 4.**
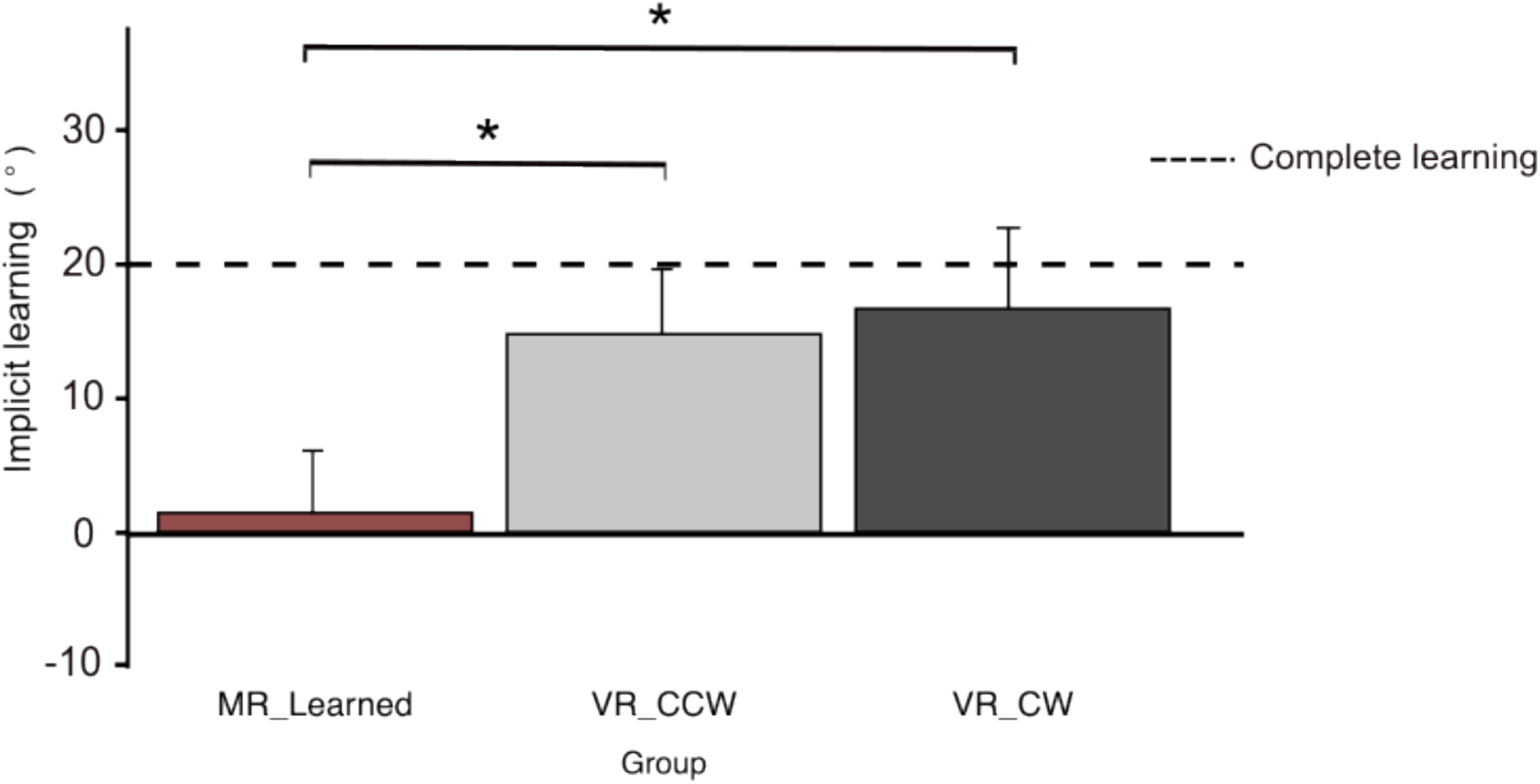
Mean implicit learning for all groups. Dashed line shows complete implicit learning. Maroon depicts the MR group, grey depicts the VR-CCW group and black depicts the VR-CW group. Error bars reflect standard error of the mean. Asterisks denote group comparisons where differences were statistically significant (*p* < 0.05).

Analysis of AE variability revealed significant main effects of group (*F*(2,30) = 73.067, *p* < 0.001, *η²* = 0.512) and time (*F*(2,30) = 8.849, *p* = 0.006, *η²* = 0.059). Participants in the MR group had more variable AEs than participants in the VR-CW group (*p* < 0.001) and VR-CCW group (*p* < 0.001). Across all participants, AE variability was significantly lower in late learning trials compared to early learning trials (*p* = 0.006; Figure 3C). AE variability did not differ across targets (*F*(1,30) = 3.338, *p* = 0.076, *η²* = 0.006).

### Implicit Learning

Analysis of implicit learning revealed a main effect of group (*F*(2,30) = 24.832, *p* < 0.001, *η2* = 0.623). Post hoc analysis indicated that the extent of implicit learning was significantly less for the MR group (M = 1.4°, SD = 8.6°) compared to the VR-CW group (M = 16.7°, SD = 2.7°, *p* < 0.001) and VR-CCW group (M = 14.8°, SD = 3.2°, *p* < 0.001). The magnitude of implicit learning in the VR-CW group was not different from VR-CCW group (*p* = 0.702).

In the MR group, implicit learning was not significantly different from zero (*t*(10) = 0.559, *p* = 0.6), and Bayesian analysis provided support for the null hypothesis (*BF₁₀* = 0.340) suggesting no true difference between implicit learning and zero in the MR group. In contrast, VR-CW and VR-CCW groups showed significant implicit learning (VR-CW: *t*(10) = 15.1, *p* < 0.001; *BF₁₀* = 5.445 x 10^6^; VR-CCW: *t*(10) = 20.8, *p* < 0.001; *BF₁₀* = 338426.099), indicating strong evidence in favor of a true effect.

## Discussion

This study investigated the contribution of implicit processes to learning a small MR distortion. The contribution of implicit processes was directly compared between MR and VR learning. Only participants who had learned the MR or VR distortion were included in our analyses. All participants in the VR groups learned their respective distortions and 11 out of 20 MR participants learned the MR distortion (see Supplementary File for data of MR participants that did not learn). Although the extent of learning by the end of the learning block was not reliably different between MR and VR groups, the magnitude of implicit learning differed significantly between VR and MR groups. Specifically, the MR group did not show evidence of implicit learning, while implicit processes contibruted to learning for both the VR-CW and VR-CCW groups. Further, we also found that reaction time (RT) was longer for participants in the MR group compared to both VR groups. These findings indicate that, unlike VR learning, MR learning does not rely on implicit processes, further supporting the view that MR engages distinct learning processes from VR learning.

To assess implicit learning, we measured persistent reaching adjustments during no-cursor exclusion trials (i.e., aftereffects), in which participants were instructed to aim directly at the target without applying what they had learned. This method is part of the Process Dissociation Procedure (Werner et al., 2015), which has been used to measure implicit processes in visuomotor learning studies (see Heirani Moghaddam et al., 2021, 2022; Modchalingam et al., 2019; Neville & Cressman, 2018; Werner et al., 2015, 2019). In the current study, the VR groups demonstrated robust aftereffects of approximately 15°, which confirmed that implicit processes contributed to VR learning. In contrast, the absence of such aftereffects in the MR group, despite successful reach adjustments by the end of the learning block, indicated that MR learners did not engage implicit processes when learning the MR distortion.

In agreement with our results, prior research in VR learning has shown that implicit processes primarily drive learning when reaching with a small VR distortion (e.g., distortion is less than 30°; Henriques & Cressman, 2012; Modchalingam et al., 2019; Neville & Cressman, 2018; Werner et al., 2015). Specifically, the magnitude of aftereffects observed in the VR groups in the present study is consistent with previous reports (see Bond & Taylor, 2015; Krakauer et al., 2005; Taylor et al., 2014). These implicit processes are proposed to be driven by a sensory prediction error, which arises due to the discrepancy between expected and actual sensory consequences of the movement (Wolpert & Kawato, 1998; Henriques et al., 2013). When the distortion exceeds 30°, explicit (i.e., cognitive strategy) contributions have also been shown to play a role in VR learning (14,15,21,23).

The lack of implicit learning in our MR group aligns with previous research that employed larger distortions (see Gastrock et al., 2024; Wang & Taylor, 2021; Wilterson & Taylor, 2021). Prior research has typically involved large MR distortions (i.e., 45° or greater), where explicit processes would be expected to dominate, in accordance with the VR literature (13–15,23). To directly test whether a smaller MR distortion might engage implicit learning, this study used a small, 20°, MR distortion. Despite the reduced distortion size, our results were consistent with studies using larger MR perturbations (6–8). That is, participants successfully adjusted their reaches during the learning block but demonstrated no implicit learning when visual feedback was removed.

In the current study, participants in the MR group exhibited significantly longer RT compared to those in the VR groups. Previous work has shown that increased RTs are often associated with the engagement of conscious, strategic mechanisms in motor learning (Fernandes et al., 2021; Haith et al., 2015; McDougle & Taylor, 2019; Taylor et al., 2014). For example,

Haith et al. (2015) examined VR learning with either long or restricted preparation times using a large (45°) rotation. They found that learning was significantly more effective when preparation time was long, as participants could engage in explicit strategies, whereas restricted preparation times limited learning to implicit processes. For small rotations, where explicit contributions are minimal, one would expect little to no effect of preparation time. Gastrock et al. (2024) and Telgen et al. (2014) recently reported elevated RTs in MR learning compared to VR learning. The RT differences, in addition to the lack of implicit adaptation observed in our study, provide behavioral evidence that MR learning engages distinct processes from VR learning.

The absence of implicit learning in MR tasks reflects differences in how MR and VR distortions are learned. As suggested by Gastrock et al. (2024) and Telgen et al. (2014), MR learning may require the formation of an entirely new sensorimotor mapping (e.g., control policy), rather than a modification of an existing sensorimotor mapping, regardless of MR distortion size. In the current study, participants in the MR group exhibited greater movement variability across both early and late learning compared to participants in the VR group. This increased variability may reflect an evolving or unstable sensorimotor mapping as participants attempted to construct a novel control policy to compensate for the mirrored cursor feedback.

Dhawale, Smith, and Ölveczky (2017) recently proposed that motor variability can serve as a mechanism for exploration, allowing the motor system to sample different movement strategies in response to novel or uncertain task demands. Similarly, Will and Stenner (2024) emphasize that variability can reflect the learner’s engagement with the broader workspace, particularly when the task requires reorganization of existing motor plans. For example, in the MR task used here, participants were required to reorganize their motor plan by moving their hand in the opposite direction of the target in order to guide the cursor accurately. In this context, the variability observed in MR learners may reflect purposeful exploration within a newly emerging control policy, rather than instability alone. Shadmehr, Huang, and Ahmed (2016) support this notion by showing that movement variability is often elevated early in learning, when the motor system is actively searching for effective strategies. Thus, the increased variability in MR may be a signature of the motor system’s attempt to explore (29–31) and establish a new sensorimotor map under conditions where implicit learning mechanisms are not engaged.

Despite the potential for purposeful exploration, not all participants in the MR group were able to learn the distortion. A large proportion of individuals in the MR condition (n = 9 of 20 participants) did not learn to reach in the MR environment (see Supplementary File). This is in contrast to participants in the VR group, who all learned to reach with rotated cursor feedback. The presence of non-learners exclusively in the MR group suggests that the MR task was more difficult to learn compared to the VR task. Task difficulty may be enhanced in the MR environment given the dynamic and non-uniform nature of errors experienced. The magnitude and direction of the distortion varies depending on movement execution on a trial to trial basis (8,32,33). In contrast to VR, where the rotation introduces a consistent angular offset, a MR distortion reverses the cursor’s position across the mirroring axis (y-axis) resulting in a transformation that can vary both in magnitude and direction depending on the hand’s spatial location. For example, a small deviation to the left of body midline produces a mirrored cursor movement to the right, but the extent of this reversal depends on how far the hand is from the axis. This means that the error signal changes from trial to trial, even for similar movement intentions, increasing the difficulty for learners to form a stable control policy (or sensorimotor map).

In addition to the extent of the error signal varying from trial to trial, the direction of the error produced also varies, depending on reaching direction relative to the target displayed. Prior work has shown that when error direction fluctuates, overall learning is reduced (33) and implicit adaptation is especially diminished (34). One explanation is that the motor system integrates a running history of experienced errors, weighting new ones relative to this record (32,35). When reaching with MR distortions, participants whose movement patterns generated frequent sign reversals would thus be more likely to suppress implicit learning, shifting reliance toward explicit, strategy-based learning (32). The presence of non-learners exclusively in the MR group therefore not only underscores the difficulty of the MR transformation but also provides further support for the distinction between MR and VR learning.

The current findings reinforce the proposal that MR and VR learning reflect distinct forms of motor learning. As outlined by Krakauer et al. (2019), motor learning encompasses both skill acquisition and skill maintenance, which correspond to de novo learning and motor adaptation, respectively (see also Gastrock et al., 2024). VR learning, particularly with small distortions, is characterized as a form of motor adaptation driven primarily by implicit processes, as evidenced by robust aftereffects once the perturbation is removed. In contrast, MR learning aligns with skill acquisition, involving the construction of a new sensorimotor mapping perhaps through more strategic processes. Our data support this distinction.

In summary, our findings demonstrate that implicit learning supports VR learning but does not contribute to MR learning, even under conditions designed to optimize implicit contributions (i.e., a small distortion size). MR learners exhibited greater movement variability, and a subset of participants did not learn the distortion at all, suggesting that reaching with a MR distortion presents additional challenges compared to reaching with a VR distortion (e.g., complex and dynamic distortion). The increased variability observed in participants in the MR group aligns with prior research showing higher variability when reaching with a MR distortion (28). Such variability, along with the presence of non-learners, suggests that MR learning poses unique challenges that may limit the effectiveness of implicit recalibration. Instead, successful performance in MR environments may require greater engagement of explicit processes. Future research is needed to determine which strategies best support MR learning and how they are implemented to improve performance.

## Supplementary File Figure

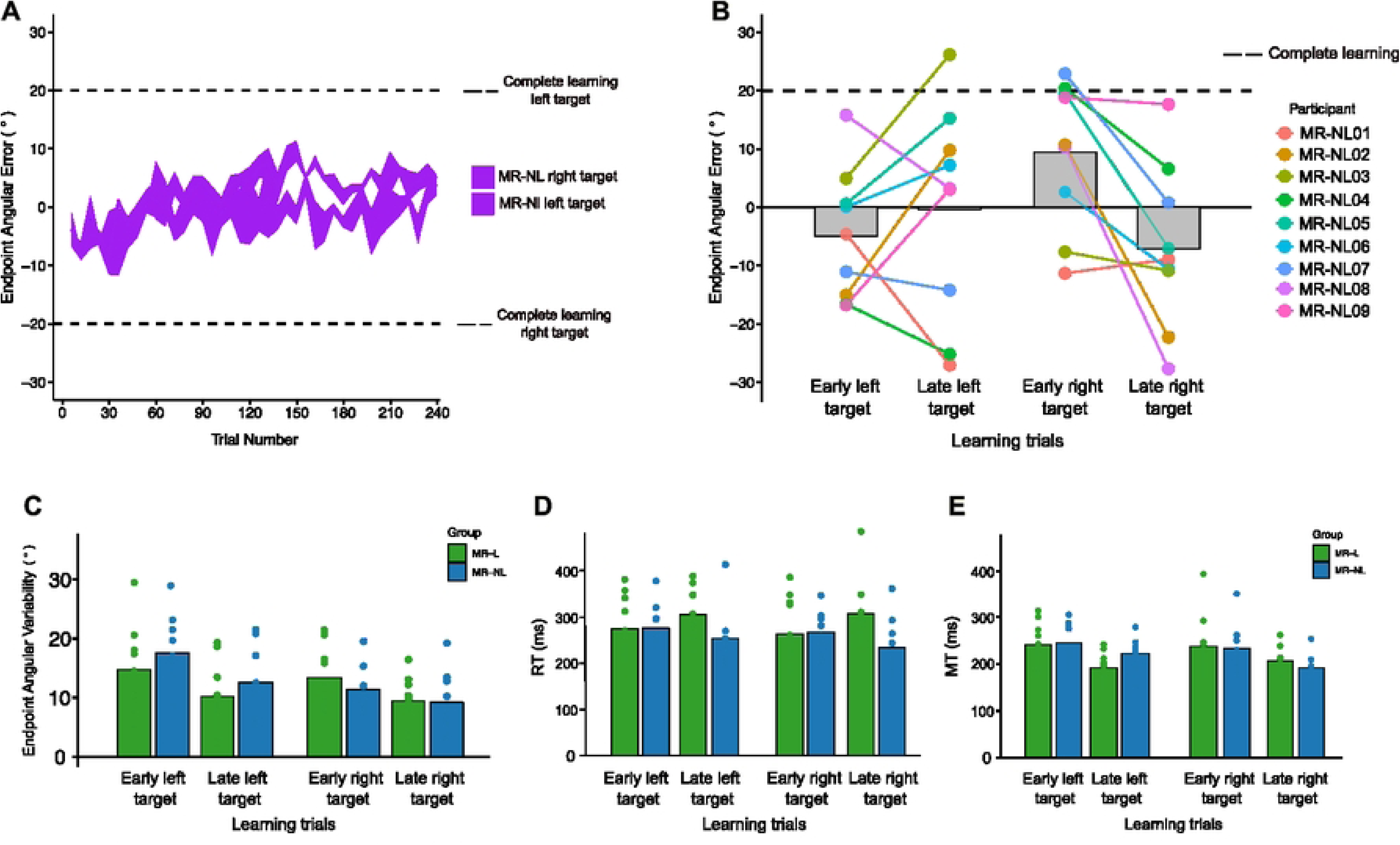

